# Click by Click Microporous Annealed Particle (MAP) Scaffolds

**DOI:** 10.1101/765529

**Authors:** Nicole J. Darling, Weixian Xi, Elias Sideris, Alexa Anderson, Cassie Pong, S. Thomas Carmichael, Tatiana Segura

**Author notes:** Corresponding author: Prof. T. Segura, Department of Biomedical Engineering, Duke University, 101 Science Drive Campus Box 90281, Durham NC 27708-0281, United States.

## Abstract

Macroporous scaffolds are being increasingly used in regenerative medicine and tissue repair. While our recently developed microporous annealed particle (MAP) scaffolds have overcome issues with injectability and *in situ* hydrogel formation, limitations with respect to tunability to be able to manipulate hydrogel strength and rigidity for broad applications still exist. To address these key issues, here we synthesized hydrogel microparticles (HMPs) of hyaluronic acid (HA) using the thiol-norbornene click reaction and then subsequently annealed HMPs into a porous scaffold using the tetrazine-norbornene click reaction. This assembly method allowed for straightforward tuning of bulk scaffold rigidity by varying the tetrazine to norbornene ratio, with increasing tetrazine resulting in increasing scaffold storage modulus, Young’s modulus, and maximum stress. These changes were independent of void fraction. Further incorporation of human dermal fibroblasts (HDFs) throughout the porous scaffold revealed the biocompatibility of this annealing strategy as well as differences in proliferation and cell-occupied volume. Finally, injection of porous HA-Tet MAP scaffolds into an ischemic stroke model showed this chemistry is biocompatible *in vivo* with reduced levels of inflammation and astrogliosis as previously demonstrated for other crosslinking chemistries.

---

Granular hydrogels are smaterials generated from hydrogel microparticle (HMP) building blocks. A solution of HMPs can form a stable bulk gel when the HMPs are above a packing density of 0.58 and jamming can occur. In our laboratory, we crosslink HMPs to each other to generate a stable scaffold, rather than rely on jamming such that the void spaces surrounding the beads are sufficiently large for cellular growth and the scaffold is structurally stable. We termed these bulk hydrogels microporous annealed particle (MAP) scaffolds due to the micron sized voids (pores) formed in between packed beads. MAP scaffolds supportmodifications with bioactive ligands to modulate cell behavior, can be used for cell culture *in vitro* ^1-8^, and support tissue ingrowth *in vivo* ^2-5^. Of particular interest is that injection of MAP scaffolds into wound sites results in reduced inflammation and improved tissue regeneration ^3,4,9,10^. Herein, we demonstrate the generation of hyaluronic acid HMPs, the modification of HMPs with ligands, and the crosslinking of HMPs to form MAP scaffolds using thiol-norbornene and tetrazine-norbornene click reactions and their use *in vitro* and *in vivo*.

Our current method to fabricate MAP scaffolds for use *in vivo* utilizes the coagulation enzyme FXIIIa, which is a transglutaminase ^2,5,11^. The HMPs are packed and linked between K and Q peptides, recognized by FXIIIa, on their surface. Because the reaction is relatively inefficient, excess reagents are used which results in poor control of crosslinking density between HMPs. Further, since the reaction is enzymatic, bond formation catalyzed by the enzyme can vary depending on environmental factors and enzyme activity. Thus, HMP linking to form a MAP scaffold cannot be easily tuned to modulate the strength and rigidity of the final scaffold. FXIIIa-crosslinked MAP typically has low storage modulus ^2^ and Young’s modulus ^12^. Biorthogonal click reactions pose as an attractive alternative to the existing annealing mechanism and have been increasingly utilized for hydrogel synthesis and functionalization for 3D cell culture and tissue engineering applications.^13-15^ Here we explored a method to make MAP building blocks by using a thiol-norbornene click reaction to form HMPs and tetrazine-norbornene click reactions to anneal them to form Tet-MAP scaffolds.

## Synthesis of Tet-MAP scaffolds using click by click reactions

Hyaluronic acid (HA) was modified to contain norbornene (HA-NB) groups through its carboxylic acid groups (**Fig. 1A** and materials and methods). NMR analysis shows that HA had 44% of its units modified with norbornene (**Sup. Fig. 1**). To generate the crosslinker, thiol-terminated polyethylene glycol (PEG) was modified with a maleimide-tetrazine bifunctional crosslinker (**Fig. 1B** and materials and methods,). NMR analysis shows 100% modification of PEG with tetrazine (**Sup. Fig. 2**). **Fig. 1C** introduces the two orthogonal click reactions that were used to generate HA-NB HMPs and Tet-MAP scaffolds (**Fig. 1D**). HMPs were generated using Thiol-ene Click chemistry occurring in an inverse suspension polymerization (**Fig. 2A**). Aqueous precursor solutions (HA-NB, dithriothreitol (DTT), Lithium phenyl-(2,4,6-trimethylbenzoyl)phosphinate (LAP)) were prepared and transferred to a continuous phase of hexane ^16^ with 3% Span-80 under nitrogen purge. In order to control microgel size, the solution was exposed to low shear through gentle pipetting and then maintained spinning at 600 rpm while UV light at 20 mW intensity initiated polymerization of the HMPs for 10 min. Particles were subsequently washed with hexane, swelled in 1% Pluronic in PBS, filtered through nylon mesh filters to gate various HMP ranges, and transitioned to PBS. The size distribution of the HMPs from each filter was characterized using an imageJ particle analyzer (**Fig. 2B**) and shown to have a mean diameter of 207.9 ± 48.21, 125.8 ± 35.51, and 76.59 ± 20.39 µm after passing through 200 µm, 100 µm, and 60 µm filters, respectively. As expected, HMP size distribution tightened as HMPs became further refined through sequential passes of decreasing filter size. The final HMPs have an HA-NB backbone crosslinked through a DTT nondegradable crosslinker.

**Figure 1.**
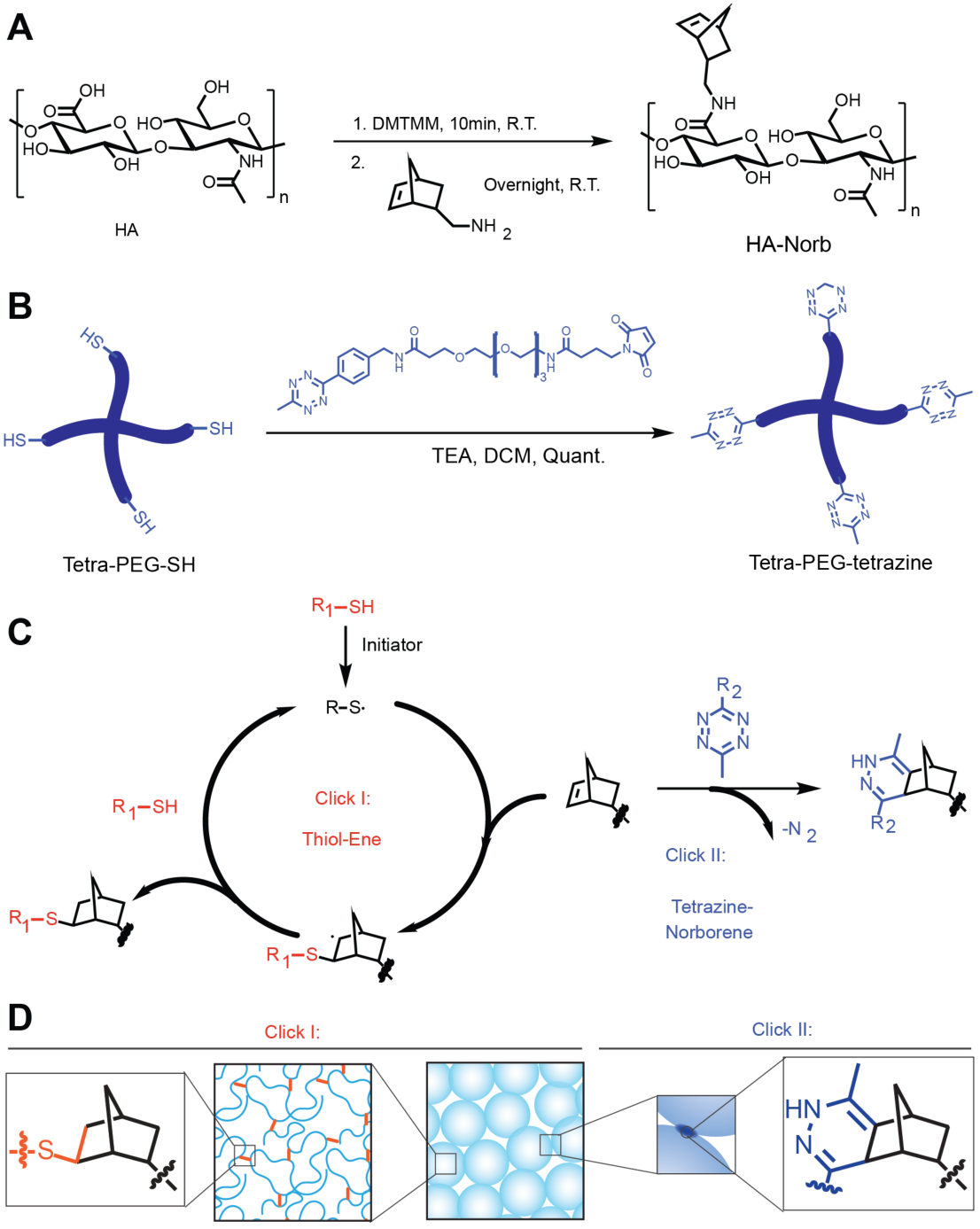
Schematic overview for the HA backbone modification, HMP crosslinker synthesis, and the click by click chemistry for both generating and annealing HMPs. (A) Hyaluronic acid-norbornene (HA-NB) was synthesized through the activation and subsequent functionalization of the HA carboxylic acid group. (B) Tetra-polyethylene glycol-tetrazine (Tetra-PEG-Tet) was synthesized through a base-catalyzed thiol-Michael addition for the purpose of annealing beads. (C) Click I is described as the radical-mediated step-growth thiol-norbornene photo-click reaction, and Click II is the tetrazine-norbornene cycloaddition click reaction. Graphical depictions of the click by click chemistry are shown in (D)for forming HMPs using thiol-norbornene (Click I) as well as annealing HMPs using tetrazine-norbornene click chemistry (Click II) to generate MAP scaffolds.

**Figure 2.**
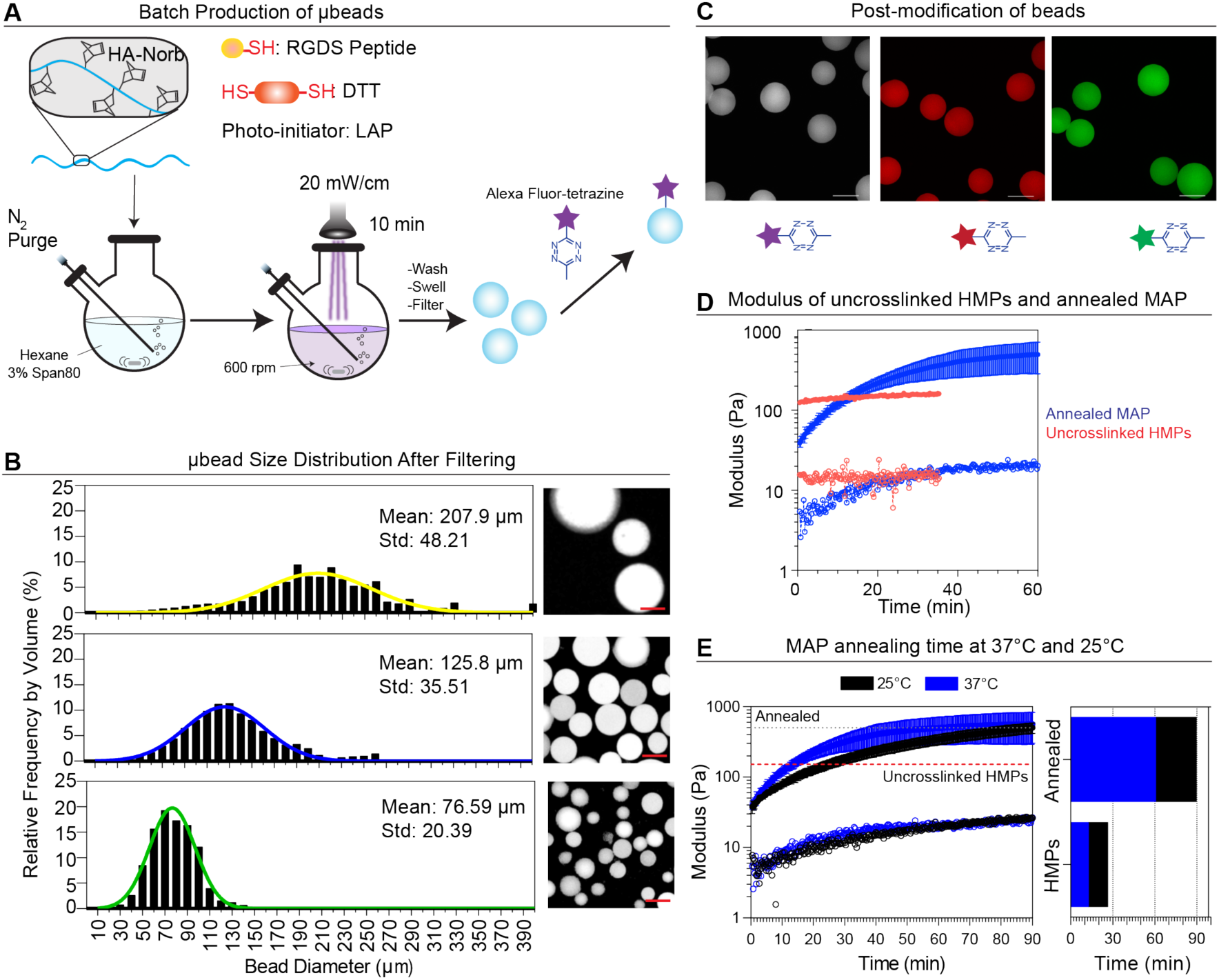
Synthesis and characterization of µbead building blocks and annealing of Tet-MAP scaffolds. (A) Overview schematic of the µbead fabrication and purification. Final bead preparations were sequentially filtered through decreasing size filters to control for µbead size. A representative image of Alexa FluorTetrazine fluorescently labeled HMPs within each filtered range is depicted. Scale bars = 100 µm. (B) Relative frequency by volume of µbead diameter distribution shows mean diameter of 207.9 ± 48.21µm, 125.8 ± 35.51µm, and 76.59 ± 20.39µm for the 200-100 µm, 100-60 µm, and 60-20 µm filter ranges. Greater than or equal to 1000 HMPs were analyzed in each range. All HMP used were from the 100-60 µm filter range. (C) Fluorescent labeling can be performed after HMP formation by post-modifying with Alexa Fluor-Tetrazine (D) Time sweep of Tet-MAP scaffold with Tet/HA ratio of 10 (blue) compared to uncrosslinked HMPs (red). (E) Additionally, Tetra-PEG-Tet was added to the packed HMPs and the G’ of Tet-MAP at 37°C (blue) and 25°C (red) was monitored over time. The red dashed line represents the G’ of packed HMPs without crosslinker. The grey dashed line represents G’ at plateau once the HMPs have annealed to form the MAP scaffold. The Tet-MAP scaffold crossed the uncrosslinked HMPs line at 12.6 min and 26.4 min and the Annealed line at 60.5 min and 89.6 min, respective to each temperature.

Particles were synthesized with excess norbornene in the HA backbone to maximize the degree of functionalization for post-fabrication modification and annealing using tetrazine-functionalized small molecules and polymers. To model small molecule functionalization and render the HMPs fluorescent, tetrazine-modified Alexa Fluor molecules were synthesized (Tetrazine-Alexa488, Tetrazine-Alexa555, Tetrazine-Alexa647, **Figure S3**) and conjugated to HMPs post-fabrication (**Fig. 2C**). In this fashion, HMPs can be functionalized with different ligands post-HMP synthesis, which enables the generation of more sophisticated environments.

HA-NB HMP annealing to form MAP scaffolds was achieved using Tetra-PEG-Tet crosslinker (**Fig. 1B, Figure S2**). As expected, unannealed jammed HMPs display typical elastic hydrogel behavior with the storage modulus being larger than the loss modulus (G’ > G”) (**Figure 2D**). Upon crosslinker addition, the storage modulus drops due to the addition of liquid resulting in the HMP being less jammed (**Fig. 2D**). However, after crosslinking, the storage modulus of the scaffold increases beyond that of the unannealed HMPs (**Fig. 2D**). The kinetics of the crosslinking reaction were explored measuring G’ over time at 1 Hz with 1% strain. G’ increases with time for all the crosslinking ratios tested reaching a plateau. Annealing was shown to be dependent on temperature, requiring 60.5 min at 37°C to reach the plateau and 89.6 min at 25°C to reach the plateau (**Fig. 2E**). To investigate the effect of the number of links between HMPs on mechanical properties of the Tet-MAP scaffold, the degree of annealing was modulated by changing the moles of Tetrazine to moles of HA ratio (Tet/HA ratio) and analyzed via shear rheology and compression, to arrive at storage and Young’s modulus, respectively. Increasing the crosslinker molar ratio increased the storage modulus of the gels from 278 Pa to 2016 Pa (**Fig. 3A**). Under compression, the modulus of the scaffold also increased with increasing Tet/HA ratio (**Fig. 2B, C**); in addition, the scaffold was able withstand more load with increasing Tet/HA ratio (**Fig. 2C**, right). Interestingly, under compression, jammed beads were unable to display elastic gel behavior with a flat stress strain curve (**Fig. 2B**, gray plot). These data show that tetrazine-norbornene chemistry allows for the tuning of the number of links between HMPs independent of HMP fabrication. Compared to our previous work with enzymatically annealed beads ^12^, tetra-PEG-Tet annealed MAP scaffolds display higher moduli (**Fig. 2C**, dash line), indicating stronger kinetics and more available crosslinking sites. Given that HMP could be further crosslinked by the Tetra-PEG-Tet, resulting in increased modulus of the HMP itself, we wanted to rule out that this alone could explain the differences observed in mechanical properties. We found that Tetra-PEG-Tet evenly diffuses approximately 20-25 µm into each bead and crosslinks it further for all of the Tet/HA ratios tested while the scaffold is annealing, (visualized by PEG-3Tet-Alexa555, **Fig. S4**). Given that the distance of crosslinking and intensity profile is similar for all conditions, the observed increase in bulk modulus for higher Tet/HA ratios is more likely a result of tighter connections between neighboring HMPs than the stiffening of individual beads.

**Figure 3.**
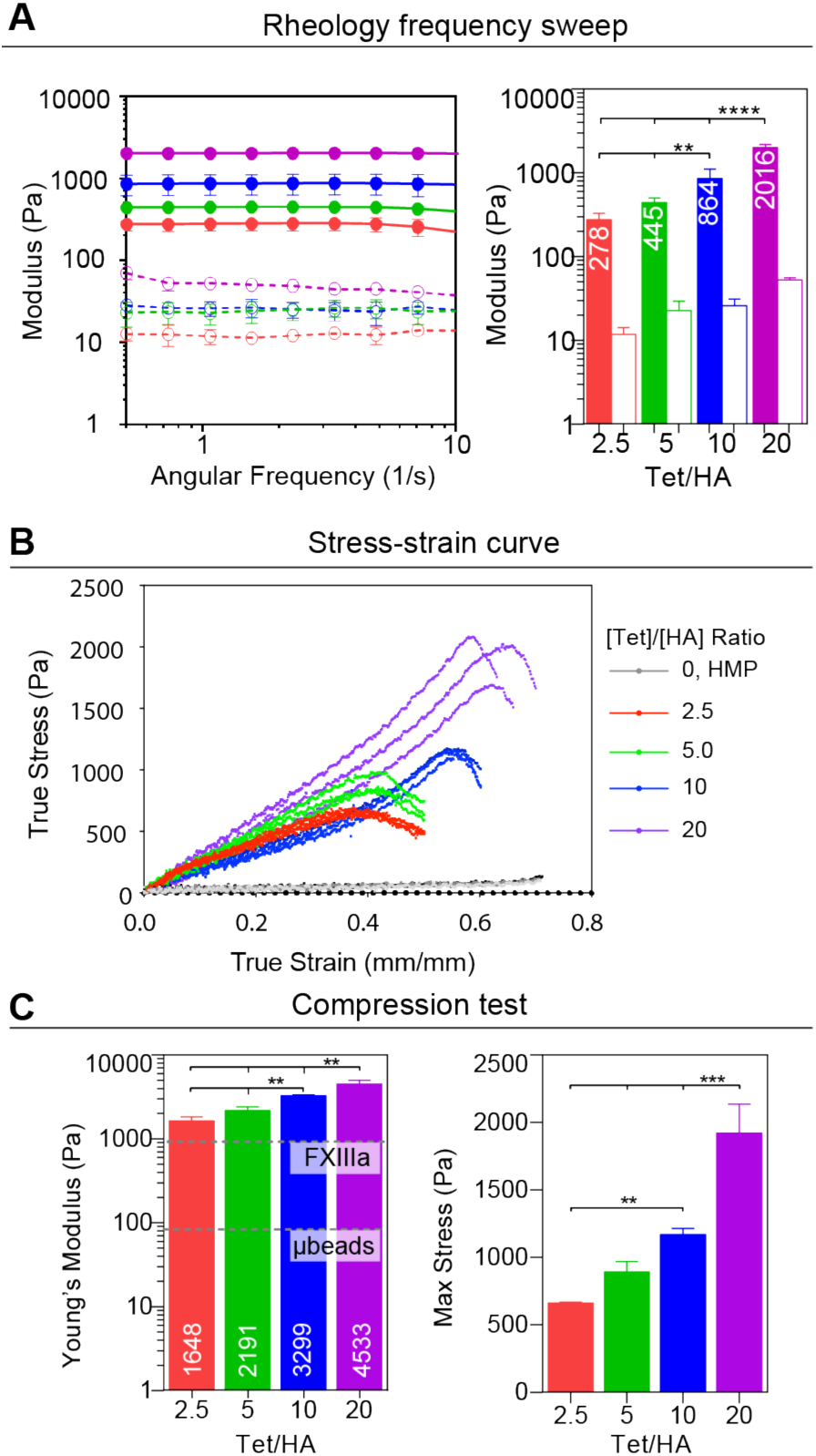
Characterization of Tet-MAP scaffolds with 2.5, 5, 10, and 20 Tet/HA ratios. (A) Frequency sweeps describing storage (solid circles/bars) and loss (open circles/bars) modulus. (B) Stress-strain curve for Tet-MAP scaffolds compared to packed HMPs. (C) Young’s modulus (FXIIIa line represents MAP annealed using previously reported K peptide, Q peptide and FXIIIa; µbead line represents packed HMP only) and max stress for each Tet/HA ratio. of Tet-MAP scaffoldsTukey’s multiple comparison test (P < 0.05). (n = 3 unless specified otherwise)

To ensure tetrazine-norbornene annealed HMPs generated MAP scaffolds that remained porous, MAP scaffolds were labeled with 2,000 kDa fluorescein-labeled dextran solution to visualize pores (**Fig. 4A**). The dextran readily diffused throughout open pores but did not penetrate HMPs. IMARIS was used to quantify overall void space and determine there were no significant differences in void fraction across the various Tet/HA ratios (**Fig. 4B**), further indicating that the mechanical changes that were observed are dependent on the interconnections between HMPs.

**Figure 4.**
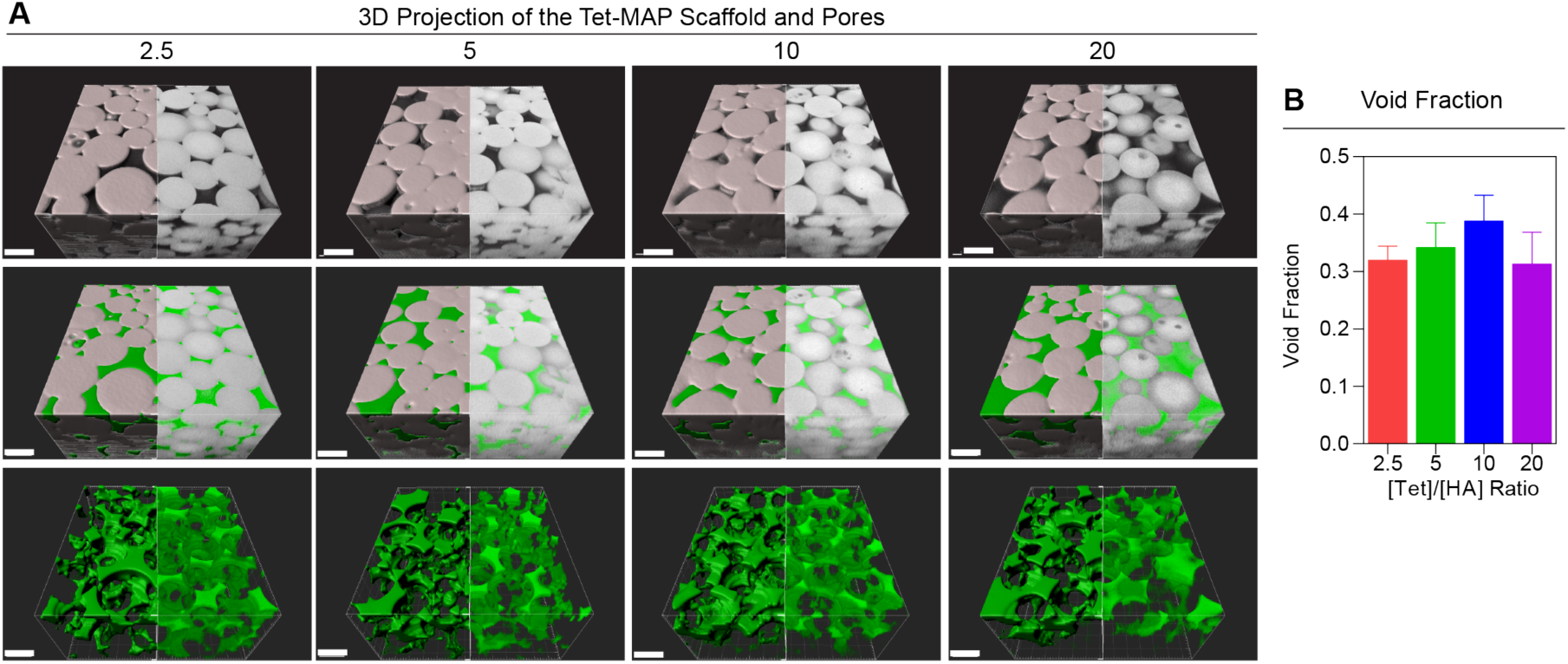
Representative images of Tet-MAP scaffolds with 2.5, 5, 10, 20 Tet/HA annealing ratios. (A) IMARIS was used to generate renderings of 300 µm z-stacks. The model generated from raw data is depicted on the left half of each image and the raw data is depicted on the right half of each image. µbeads are shown in white and the void (filled with 2,000 kDa FITC-dextran) is shown in green; scale bars = 200µm. (B) Void fraction (computed using IMARIS) of Tet-MAP scaffolds with 2.5, 5, 10, and 20 Tet/HA ratios. Tukey’s multiple comparison test (P < 0.05). (n = 3 unless specified otherwise)

## Cell culture within tetrazine-norbornene annealed HA-MAP scaffolds

Given that the tetrazine-norbornene reaction results in the generation of one molecule of nitrogen, we wanted to ensure that this chemistry is biocompatible *in vitro* and *in vivo*. RGD-modified hyaluronic acid was used to generate HMPs. Following our previous protocols for culturing cells in MAP scaffolds, HMPs and cells (human dermal fibroblasts (HDFs)) were mixed together along with the Tetra-PEG-Tet crosslinker and allowed to anneal at 37°C. The kinetics of the crosslinker reaction are slow enough to allow sufficient mixing of the cells and HMPs to ensure even seeding. Overall cell viability was assessed with the lowest and highest annealing ratios used, 2.5 and 20, and showed high survival, >95%, for both Tet/HA annealing ratios at 1 day and 8-days post seeding (**Fig 5A-C**). We further assess biocompatibility using long term culture with 2.5 Tet/HA annealing ratio. HDFs showed significant proliferation over time (**Fig. 5C**) and degree of cell spreading (**Fig. 5B**), showing that the chemistry does not negatively impact cell growth long term. The number of cells increased by approximately3-fold at two weeks relative to the average of all day 2 nuclei counts (**Fig. 5D**). HDF cell spreading was assessed at 2, 5, and 14-days via confocal microscopy. Cells were fixed and stained for F-actin. Both the degree of spreading and number of cells increased with time (**Fig. 5E**). By day 2, significant cell spreading was observed, which occurred within the void space of the MAP scaffold (**Fig. 5F**). IMARIS 3D rendering of the cells over time (**Fig. 5G**) as well as the cells and scaffold (**Fig. 5H**) show again that in the volume of the scaffold, the cells reside in the interstitial space and that the cell number increases over time.

**Figure 5.**
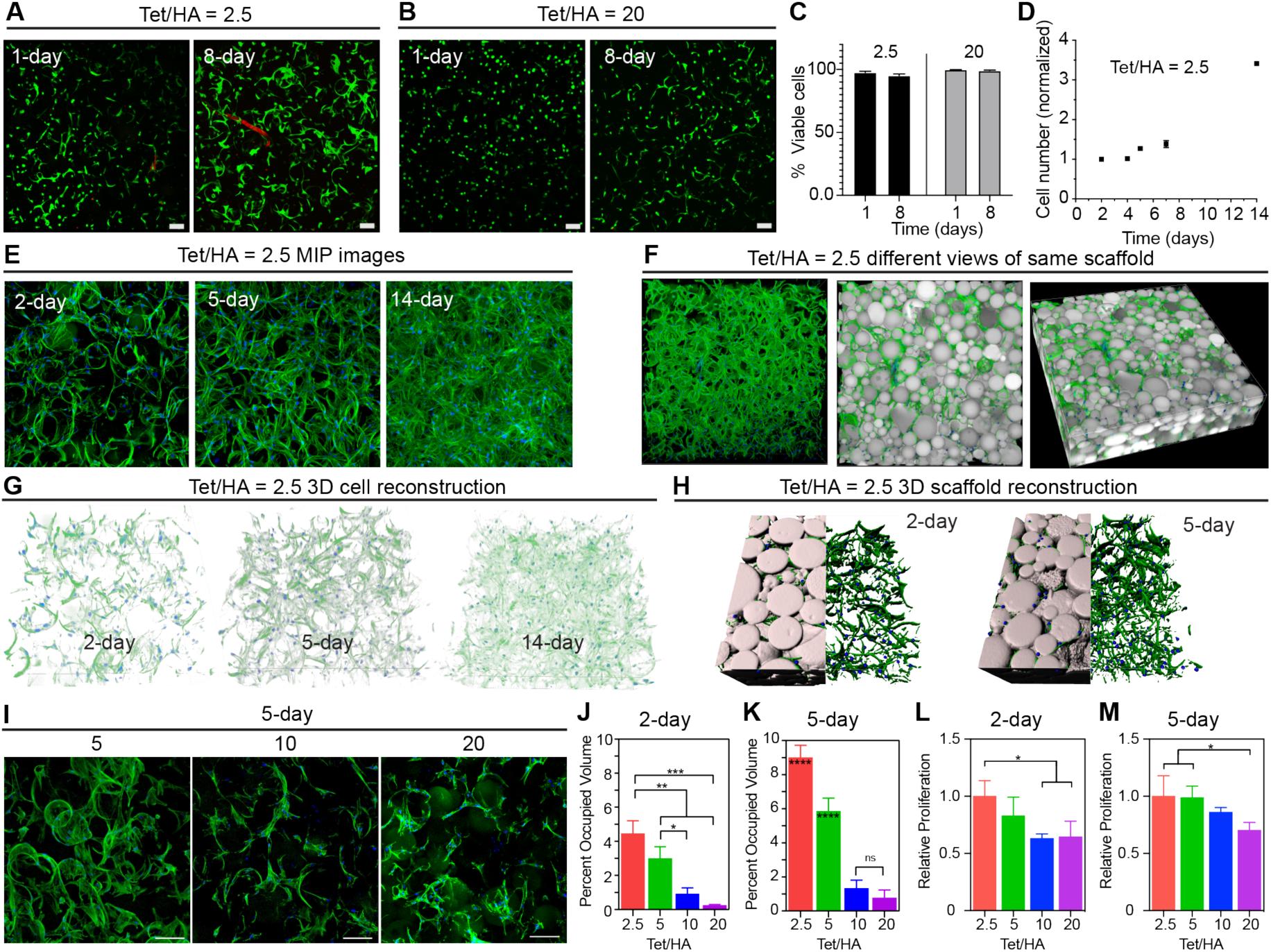
*In vitro* cellular response to Tet-MAP scaffolds. (A) Representative images and analysis of HDFs (F-actin labeled green) in Tet-MAP scaffolds 1 and 8-days post-encapsulation with Tet/HA annealing ratios of (A) 2.5 and (B) 20. (C) Overall cell viability was assessed with the lowest and highest annealing ratios used, 2.5 and 20, and showed high survival (>95%) for both Tet/HA annealing ratios at 1 day and 8-days post-encapsulation. (D) The number of cells relative to the average of all day 2 nuclei counts. (E) Representative maximum intensity projection (MIP) images of HDFs in 2.5 Tet/HA scaffolds 1 and 8-days post-encapsulation. (F) 3D projection of the 300 µm z-stacks with HMPs shown in white, F-actin shown in green, and cell nuclei (labeled with DAPI) are shown in blue. (G) IMARIS analysis of HDF nuclei in z-stacks to obtain information about cell number and proliferation (H) IMARIS renderings of confocal microscopy z-stacks. The model generated from raw data is depicted on the left half of each image and the raw data is depicted on the right half of each image. (I) Cells were able to spread in all the scaffolds as determined by phalloidin staining and confocal microscopy imaging. The percent volume occupied by HDFs F-actin measured by IMARIS at three different regions in a scaffold of each Tet/HA on (J) 2 days and (K) 5 days post-encapsulation. Relative proliferation measured by PrestoBlue at (L) 2 days and (M) 5 days. Turkey’s multiple comparison test (P < 0.05). (n = 3 unless specified otherwise)

From our viability experiment it appeared that HDFs spread more in the 2.5 versus 20 Tet/HA ratio gels. Since the viability of the scaffold is the same for both conditions, we further investigated the effect of HMP annealing degree on spreading and proliferation. Cell-loaded MAP scaffolds were generated with 2.5, 5, 10, or 20 Tet/HA ratio and analyzed for cell spreading, cell metabolic activity, and overall cell volume at 2 and 5-days post seeding. Cells were able to spread in all the scaffolds as determined by phalloidin staining and confocal microscopy imaging (**Fig. 5I**). Analysis of the occupied volume via IMARIS 3D analysis software reveals that cells seeded in 2.5 Tet/HA ratio scaffolds contain more cell volume than cells in the other Tet/HA ratio scaffolds for all the time points tested (**Fig. 5,K**). The percent of occupied cell volume increases for all Tet/HA scaffolds between days 2 and 5 post-encapsulation, with noticeable increases in cell spreading for Tet/HA ratios of 2.5 and 5. Analysis of metabolic activity at 2 and 5-days, revealed that cells seeded in 2.5 and 5 Tet/HA ratio scaffolds resulted in a higher proliferation rate compared to 10 and 20 Tet/HA (Fig. 5L,M). Taken together these data indicate that cells seeded on stiffer scaffolds generated by the 10 and 20 Tet/HA crosslinker ratio result in significantly less cell spreading and proliferation.

## In vivo biocompatibility of Tet-MAP scaffolds in brain tissue

We next assessed the biocompatibility of tetrazine-norbornene crosslinkers *in vivo*. We chose to inject the material into a brain wound environment because it is an environment with an increased population of immune cells and because of our interest in stroke repair. We have previously shown that hydrogels can be injected into the stroke core without damaging the brain or causing swelling ^5,17-19^. The peri-infarct space, region that surrounds the stroke core, is the tissue of the brain that undergoes the most substantial repair post-stroke^17^. Thus, injecting material into the stroke core itself provides an opportunity to develop regenerative tissue engineering therapies promote brain repair post-stroke. Previous work in our lab from Nih and Sideris showed that the use of HA-MAP gels annealed by FXIIIa were biocompatible and resulted in less microglia infiltration and astrocytic scarring compared to a nonporous hydrogel of the same biochemical composition ^5^. Here we aimed to assess the effect, if any, of using the tetrazine-norbornene reaction for HMP annealing to form *in situ* MAP scaffolds in the brain. In our experimental timeline, a photothrombotic stroke is induced in the motor cortex at t = 0 and five days (t = 5-days) later a gel is injected at the same coordinates where the stroke was induced (**Fig. 6A**). Five days after stroke, the stroke core is surrounded by an astrocytic scar (**Fig. 6B**). Microphage/microglia are observed both inside the stroke core and in the surrounding peri-infarct tissue (**Fig. 6B**). Following hydrogel injection, no obvious damage to the surrounding brain tissue was observed (**Fig. 6C**). Similar to FXIIIa-crosslinked HA-MAP scaffolds, we found that tetrazine-norbornene crosslinked HA-MAP (Tet-MAP) scaffolds reduced the number of inflammatory monocytes (CD11b+) cells (**Fig. 6D**). Quantification of the infarct and peri-infarct region for CD11b+ area (**Fig. 6E**) reveals that the area covered by CD11b+ cells is statistically lower for animals treated with Tet-MAP gel (**Fig. 6F**). Analysis of the astrocytic scar again revealed that similar to FXIIIa crosslinked HA-MAP scaffolds, injection of Tet-MAP into the stroke core visibly reduced astrocytic scar thickness (**Fig. 6G**). Quantification of the scar thickness (**Fig. 6H**) revealed statistical significance for thickness reduction with Tet-MAP compared to sham groups (**Fig. 6I**). Together these data show that tetrazine-norbornene chemistry can be used as a crosslinker *in vivo* without causing more inflammation or reactive astrogliosis.

**Figure 6.**
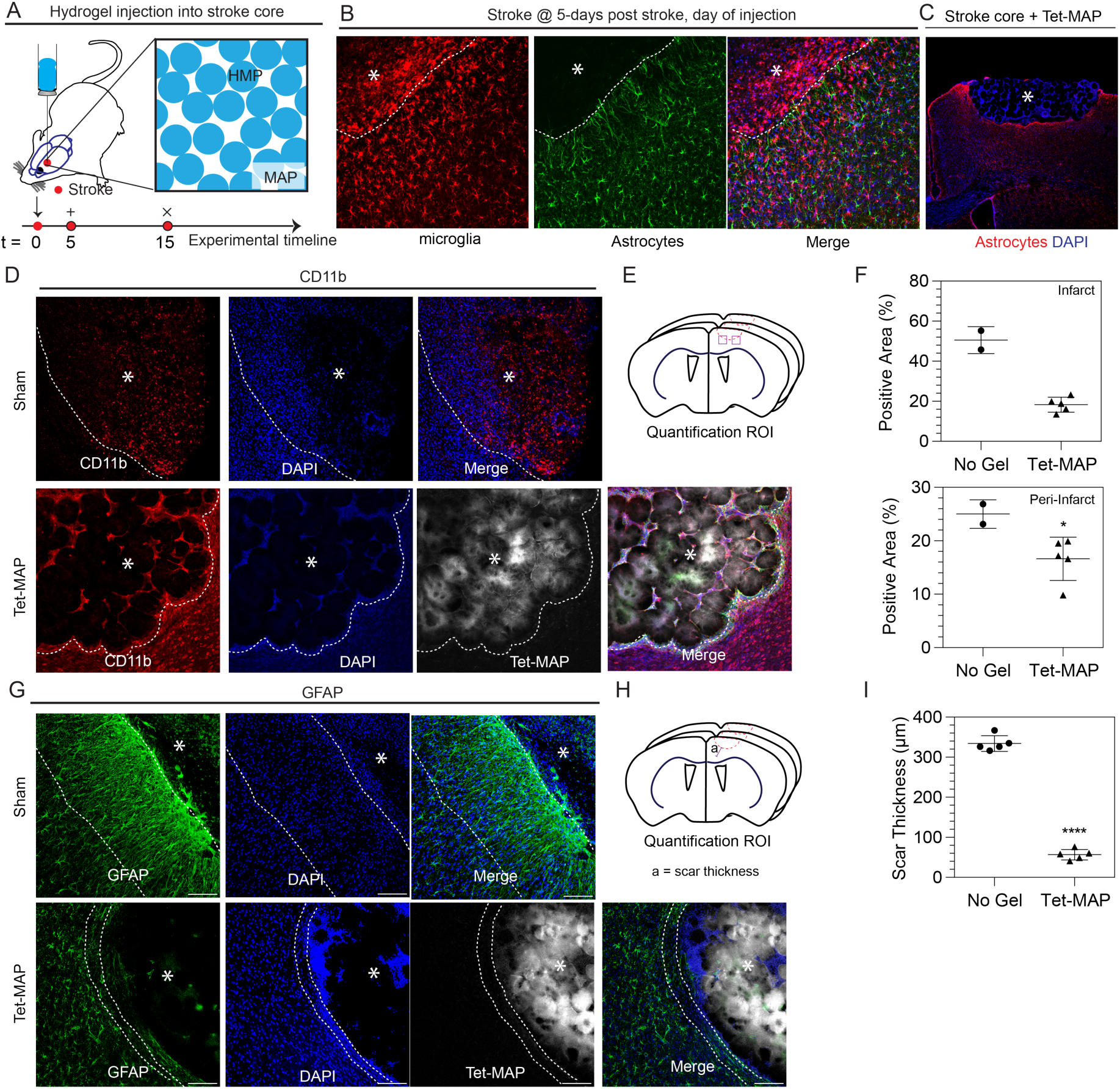
*In vivo* cellular response to Tet-MAP scaffold in a PT stroke model. (A) Overview schematic of PT stroke timeline and stroke injection concept. (B) Images showing the microglia/macrophage (IBA1) and astrocytic (GFAP) response at 5-days post stroke, which is the day of hydrogel injection. (C) Image showing brain + TetMAP at 2-weeks post stroke. The gel area is completely fills the stroke core. Fluorescent images at 2-weekw of (D) CD11b staining showing the post-stroke microglial response. (E) The ROI for quantification is highlighted in for (F) analysis of CD11b+ area in both the infarct and peri-infarct regions. (G) GFAP staining showing the post-stroke astrocyte response and (H) the ROI for quantification of GFAP+ response in terms of scar thickness (I). (E) analysis Scale bars = 100µm; white star indicates the infarct region Tukey’s multiple comparison test (P < 0.05). (n = 5 unless specified otherwise)

## Conclusion

Tet-MAP scaffolds crosslinked using click-by-click chemistry were developed to be able to easily tune material properties for the study of cell behavior *in vitro* and for therapeutic use *in vivo*. By manipulating the Tet/HA annealing ratio, we were able to tune the rigidity of the scaffold which influenced how HDFs behave within the porous scaffold. In particular, we observe that MAP scaffolds containing lower annealing degree allowed cells to proliferate faster and result in substantial cell spreading compared to MAP scaffold that contained a higher annealing degree, likely due to the ability of cells to re-arrange their local environment in a less crosslinked, less stiff scaffold. Further, we find that tetrazine-norbornene chemistry is biocompatible *in vivo* using an experimental ischemic stroke model in mice with our results mirroring what we observed for HMPs crosslinked with factor XIIIa enzyme. In particular, we found that *in situ* annealing of HMPs to form MAP scaffolds in the stroke cavity lowers the CD11b+ cell population and decreases the scar thickness surrounding the stroke. This platform holds potential to investigate how different HMP properties like size, shape, mechanical strength, topography/porosity, and biochemical composition impact cellular responses and tissue repair.

## Experimental Section

### Synthesis of Tetrazine Reactive Monomers

Tetra-polyethylene glycol-tetrazine (Tetra-PEG-Tet) was synthesized through a base-catalyzed thiol-Michael addition by dissolving 100 mg of tetra-PEG-SH (MW: 20,000 Da) (NOF America, White Plains, NY) and 15 mg of methyltetrazine-PEG4-maleimide (MW: 514.53 Da) (Kerafast, Boston, MA) in 0.5 mL CDCl_3_ (maleimide/SH ratio of 1.05), then adding 1 µL of triethylamine (TEA) (0.5 molar equivalent). (**Scheme 1B**) The mixture was stirred at room temperature for 4 hrs. The product was precipitated in 50 mL cold diethyl ether and confirmed by ^1^H-NMR with 98% conversion. (Figure S4)

Alexa Fluor 647 C2-tetrazine (Alexa647-Tet) was synthesized through two base-catalyzed thiol-Michael addition reactions in series. (Figure S3) First, by dissolving 2.8 mg of HS-PEG-SH (MW: 3500 Da) (JenKem Technology USA, Plano, TX) (1 molar equivalent to Alexa Fluor C2 maleimide) and 0.41 mg of methyltetrazine-PEG4-maleimide (MW: 514.53 Da) (Kerafast, Boston, MA) (1 molar equivalent to Alexa Fluor C2 maleimide) in 0.17 mL CDCl_3_, then adding 0.11 µL of triethylamine (TEA) (0.5 molar equivalent). The mixture was stirred at room temperature overnight. Next, by dissolving 1 mg Alexa Fluor 647 C2 maleimide (MW: 1250 Da) (Thermo Fisher Scientific) in 0.17 mL CDCl_3_, then adding the reaction mixture and 0.11 µL of triethylamine (TEA) (0.5 molar equivalent). The mixture was stirred at room temperature overnight. The product was precipitated in 10 mL of cold diethyl ether and dried under vacuum overnight. The resulting product was dissolved in dimethylformamide at 1 mg/mL and stored at -20°C.

### µbead Modification

HA-Norb HMP were modified post-fabrication via inverse electron demand Diels-Alder tetrazine-norbornene click reaction, in which excess norbornene groups on the HMP were functionalized with Alexa647-Tet to fluorescently tag the HMP. Briefly, the HMP were incubated in 1xPBS containing a final Alexa647-Tet concentration of 0.005 mM at 37°C for 1 hour under agitation (i.e. 200 µL of HMP were combined with 100 µL 1xPBS containing 0.015 mM Alexa647-Tet (1:13 dilution of the 1 mg/mL Alexa647-Tet stock)). Upon completion of the functionalization, the suspended HMP were pelleted by centrifuging at 14,000 rcf for 5 minutes. The HMP were washed three times with 1xPBS and recovered with the same centrifugation conditions. The labeled HMP were stored at 4°C until further use.

### Tet-MAP Scaffold Assembly

Macroporous annealed particle (MAP) scaffolds were assembled via inverse electron demand Diels-Alder tetrazine-norbornene click reaction, in which excess norbornene groups on the HMP were linked by tetra-PEG-Tet to form tetrazine mediated MAP (Tet-MAP) scaffolds. The theoretical HA-Norb concentration (0.18 mM) in a volume of pelleted HMP was first calculated by converting the total µbead weight percent, determined by lyophilizing 100 µL of unlabeled HMP to get the total mass, using the mass fractions of the µbead contents. This theoretical concentration was then used to determine the final tetra-PEG-Tet annealing concentrations: 0.11 mM, 0.22 mM, 0.44 mM, and 0.89 mM for tetrazine/HA-Norb (Tet/HA) ratio of 2.5, 5, 10, and 20. The volume of tetra-PEG-Tet added to the HMP was consistent across the four Tet/HA conditions, 5/6 of the final Tet-MAP scaffold volume was HMP and 1/6 of the volume was concentrated tetra-PEG-Tet (0.22 mM, 0.45 mM, 0.89 mM, and 1.79 mM, respectively). Tet-MAP scaffolds were formed by combining and mixing HMP with concentrated tetra-PEG-Tet at the desired Tet/HA ratio and incubated for 1 hour at 37°C unless stated otherwise. All Tet-MAP scaffolds were formed using the 60 - 100 µm bead size filter range.

### Animal Stroke Model, Tissue Processing, Immunohistological Staining and Image Analysis

Animal procedures were performed in accordance with the US National Institutes of Health Animal Protection Guidelines and the University of California Los Angeles Chancellor’s Animal Research Committee. The stroke model was performed as previously described. Briefly, a permanent cortical photothrombotic stroke was induced on young adult C57BL/6 male mice (8–12 weeks) obtained from Jackson laboratories (Bar Harbor, ME). The mice were anesthetized with 5% isoflurane and placed in a stereotactic setup. The mice were kept at 2.5% isoflurane in N2O:O2 for the duration of the surgery. A midline incision was made and Rose Bengal (10 mg/mL, Sigma-Aldrich) was injected intraperitoneally at 10 μL/g of mouse body weight. After 5 minutes, a 2-mm diameter cold fiberoptic light source was centered at 0 mm anterior/1.5 mm lateral left of the bregma for 18 minutes and a burr hole was drilled through the skull in the same location. All mice were given sulfamethoxazole and trimethoprim oral suspension (TMS, 303 mL TMS/250 mL H20, Amityville, NY) every 5 days for the entire length of the experiment.

Five days post-stroke, freshly mixed HMP with concentrated tetra-PEG-Tet at a Tet/HA ratio of 5 was loaded into a 25 µL Hamilton syringe (Hamilton Reno, NV) connected to a pump and 6 µL of microgels were injected into the stroke cavity using a 30 gauge needle at a depth of 0.8 mm and the same stereotaxic coordinates as above at an infusion speed of 1 µL/min. The needle was withdrawn from the mouse brain 5 min after the injection to allow for annealing of the Tet-MAP scaffold.

Ten days post-injection, mice were sacrificed via transcardial perfusion of 1xPBS followed by 40 mL of 4% PFA. The brains were isolated and post-fixed in 4% PFA overnight and submerged in 30 (w/v) % sucrose solution for 48 h. Tangential cortical sections of 30 µm thickness were sliced using a cryostat and directly mounted on gelatin-subbed glass slides. Slides not immediately stained were kept at -80°C.

Each slide was rinsed with 1xPBS for 10 minutes at room temperature, dried and outlined with a hydrophobic pen (ImmEdge Hydrophobic Barrier PAP Pen, Vector Labs). The slides were then incubated in a blocking solution containing 1xPBS, 0.3% Triton X-100 and 10% normal donkey serum for 1-2 h at room temperature. The slides were then incubated in the primary antibody at the appropriate dilution in blocking solution overnight at 4°C. After 3× 10 minute washes in 1x PBS, the slides were incubated in the secondary antibodies at the appropriate dilution in blocking solution for 2 hours at room temperature. The slides were then counterstained with the nuclear marker DAPI (1:500, Invitrogen) for 15 minutes at room temperature. After allowing the slides to dry at room temperature after 3× 10 minute washes in 1xPBS, the slides were dehydrated in ascending ethanol baths, incubated in xylene and mounted in mounting medium (DPX, Sigma-Aldrich). Primary antibodies were used as follows: rat anti-glial fibrillary acidic protein (GFAP, 1:100, Abcam, Cambridge, MA, USA) for astrocytes and rat anti-CD11b (1:100, Abcam, Cambridge, MA, USA) for microglial and macrophage cells. Secondary antibodies, matching the desired primary antibody host, conjugated to Alexa Fluor 488 (1:500, Jackson Immuno Research, West Grove, PA) were used. A Nikon-C2 laser scanning confocal microscope with a 20x air objective used to take fluorescent images represented as maximum intensity projections.

Analyses were performed on microscope images of three coronal brain levels at +0.80, −0.80, and −1.20 mm according to bregma, which consistently contained the cortical infarct area. The thickness of scar was measured on the ischemic boundary zone within the ipsilateral hemisphere on three sections stained for GFAP. The astrocytic (GFAP) infiltration into the stroke cavity was measured in ImageJ as the shortest distance from a GFAP^+^ cell to the ischemic boundary zone. The inflammation (CD11b) positive area in the infarct and peri-infarct areas were quantified using ImageJ and expressed as the area fraction of positive signal per total area (%).All measurements were averaged across sections and presented per animal.

## Supporting Information

Supporting Information is available from the Wiley Online Library or from the author.

## Acknowledgements

Author contributions: N. J. D. and W. X. contributed equally to this work. N. J. D., W. X., E. S., and T.S. designed the experiments, N. J. D., W. X., E. S., and C. P. performed experiments and analyzed the results. N. J. D., W. X., and T.S wrote the manuscript, with input from all authors. The authors would like to thank Dr. Talar Tokatlian for her assistance editing the manuscript.

## Supporting Information

### Synthesis of Norbornene Reactive Monomers

Hyaluronic acid-norbornene (HA-Norb) was synthesized through the subsiquent activation and functionalization of the HA carboxylic acid group by dissolving 1.0 g of HA (MW 60,000 Da) (Genzyme, Cambridge, MA) and 3.1 g 4-(4,6-Dimethoxy[1.3.5]triazin-2-yl)-4-methylmorpholinium chloride (DMTMM) (MW: 294.74 Da) (TCI America, Portland, OR) (4 molar equivalents) each in 40 ml of 200 mM MES buffer, pH 5.5, combining the solutions and allowing the reaction to stir for 10 min. Then 0.677 mL of 5-Norbornene-2-methanamine (a mixture of isomers) (NMA) (TCI America, Portland, OR) (2 molar equivalents) was added dropwise into the mixture. The reaction was stirred at room temperature overnight and then precipitated in 1L of 100% Ethanol. (**Scheme 1C**) All precipitates were collected and dissolved in 2M brine solution and dialyzed against DI water for 30 minutes, 1M brine solution for 30 minutes. This dialysis process was repeated 3 times and then dialyzed against DI water for 24 hours. The final solution was collected and lyophilized to yield the final product. HA-Norb was confirmed by ^1^H-NMR with 44% Norb functionalization. ^1^H NMR shifts of pendant norbornenes in D_2_0, d6.33 and d6.02 (vinyl protons, endo), and d6.26 and d6.23 ppm (vinyl protons, exo), where compared to the HA methyl group δ2.05 ppm to determine functionalization. All equivalents are based on the moles of the HA repeat unit. (Figure S1, Figure S2)

### µbead Synthesis

HA-Norb HMP were prepared via inverse suspension photo-polymerization, in which HA-Norb containing clustered Ac-RGDSPGERCG-NH_2_ (RGD) (Genscript, Piscataway, NJ) was polymerized with dithiothreitol (DTT) (Sigma-Aldrich) via radically-mediated thiol-norbornene click reaction in an aqueous phase that was suspended in an organic phase. (Figure 1A) Briefly, the organic phase was comprised of 10 mL of hexane containing 300 mg, 3 wt%, of sorbitan monooleate (Span 80) (Sigma-Aldrich). The volume of the aqueous phase was 6 mL comprised of 300 mM HEPES buffer (Thermo Fisher Scientific) at pH 8.3 with 0.5 mM of HA-Norb, 0.5 mM RGD, 3.5 mM DTT (SH/HA ratio of 14), 1.875 mM tris(2-carboxyethyl)phosphine (TCEP) (Sigma-Aldrich) (TCEP/SH ratio of 0.25) and 4.25 mM lithium phenyl(2,4,6-trimethylbenzoyl)phosphinate photo-initiator (LAP) (TCI America, Portland, OR). The RGD was initially clustered onto HA-Norb by combining 0.25 mM HA-Norb, 0.5 mM RGD (Clustering ratio of 2 RGD/HA), 1.875 mM TCEP and 0.15 mM LAP in 3.78 mL of HEPES and exposing the mixture to 10 mW/cm^2^ of UV light for 1 minutes. The mixture was then combined with the DTT and the remaining HA-Norb and LAP. This final gel precursor solution was then pipetted into a round-bottom flask containing the organic phase continuously stirring at 600 rpm and bubbled with nitrogen to minimize oxygen quenching of radicals and then mixed by pipetting up and down 9 times to generate a stable inverse suspension. The flask’s contents were then exposed to UV light at 20 mW/cm^2^ for 10 minutes to initiate polymerization.

Upon completion of the polymerization, the suspension was transferred into a conical tube and centrifuged at 1000 rcf for 1 minute and the supernatant was decanted. The HMP were washed twice with hexanes and recovered with the same centrifugation conditions. The HMP were then transferred to 10 mL 1% Pluronic F107 in PBS for 30 min to allow for swelling before sieving using 200 µm, 100 µm, 60 µm, and 20 µm (PluriSelect, Leipzig, Germany) pore size strainers. During sieving, HMP were washed with 50 mL 1xPBS. The collected HMP were then suspended in 1xPBS and washed three times by centrifuging at 14,000 rcf for 5 minutes. The HMP were then suspended in 1xPBS before autoclaving and recovered with the same centrifugation conditions. The HMP were stored at 4°C until further use.

### µbead Characterization

The size of labeled HMP were characterized by diluting labeled HMP in 1xPBS in a coverslip bottomed polydimethylsiloxane (PDMS) (Sylgard 184 PDMS, Dow Corning) well followed by acquiring a z-stacks using Nikon-C1 laser scanning confocal microscope with a 10x air objective to obtain a maximum intensity projection. These images were then analyzed using the particle analysis toolkit in ImageJ to obtain diameter measurements of ≥ 1000 HMP for each condition, which were subsequently plotted using the frequency distribution in Prism (GraphPad).

### Tet-MAP Scaffold Characterization

The kinetics of annealing Tet-MAP scaffolds were evaluated by subjecting them to oscillatory shear at 1 Hz and 1% strain were the evolution of the storage and loss moduli (i.e., G’ and G”) could be monitored until the storage modulus plateaued. The time sweep of HMP only and Tet-MAP at 25°C and 37°C were evaluated in triplicate with a plate-to-plate rheometer (Physica MCR 301, Anton Paar, Ashland, VA) using an 8 mm plate by loading 30 µL of HMP only or Tet-MAP at a 10 Tet/HA ratio between a 0.5 mm plate gap.

In addition, the swollen networks storage and loss moduli of Tet-MAP scaffolds of varying Tet/HA ratios were obtained in strain oscillatory shear at 37 °C. Three gels for each condition were prepared between two sigmacote (Sigma-Aldrich) coated glass slides (1 mm thickness spacer) and allowed to swell overnight in 1xPBS. The linear viscoelastic regime was determined through strain sweep (10 Hz; 0.05–100% strain) followed by a frequency sweep (0.01–100 Hz; 1% strain) used to determine G’ and G” at a normal force of 0.01N.

Additional mechanical testing on the Tet-MAP scaffolds was done using a 5500 series Instron. A 2.5N load cell with a 3.12 mm diameter tip was used at a compression strain rate of 1 mm/min and the hydrogel scaffold was indented 0.8 mm or 80% of its total thickness. The Young’s modulus was determined by the slope of the line in the linear region where stress is proportional to strain. The max stress or compressive strength is defined as maximum stress that a material can withstand while being compressed before breaking and was determine by the maximum point on the stress-strain curve.

The void fraction of annealed Tet-MAP scaffolds of various annealing ratios was calculate in IMARIS after a 300 µm z-stack series of the HMP and the labeled void space was acquired using Nikon-C2 laser scanning confocal microscope with a 10x air objective. The Tet-MAP scaffolds were incubated with 300 mM triethanolamine (TEOA) containing 1 µg/mL 2,000 kDa fluorescein isothiocyanate-dextran (FITC-dextran) (Sigma-Aldrich) to fill the void space in between HMP, as it is too large to penetrate the µbead’s polymer network. The z-stacks were imported into IMARIS to generate surface renders, and void space volumes were quantified as a fraction of the total volume represented by the z-stack.

### Cell Culture

Human dermal fibroblasts (HDF; Cell Applications, Inc., San Diego, CA) were encapsulated at a concentration of 2,000 cells/μL into the Tet-MAP scaffolds and cultured in Dulbecco’s modified Eagle’s medium (Thermo Fisher Scientific) containing 5% fetal bovine serum (Thermo Fisher Scientific) at 37°C and 5% CO_2_. The HMP were equilibrated in media for 30 minutes before recovering by centrifugation and annealing with one of the four concentrated tetra-PEG-Tet solutions (described above) containing 12,000 cells/μL in media. 15 µL of the Tet-MAP solutions with HDFs was pipetted into a coverglass bottomed PDMS well lined with a thin layer of 1% agarose to prevent cell attachment to the glass. After annealing, the wells were rinsed and filled with 200 µL media.

Post-encapsulation a set of gels were fixed at day 2 with 4% paraformaldehyde (PFA) (Fisher Scientific) for 15 minutes at room temperature after rinsing with 1xPBS. The cultures were permeabilized in 0.3% Triton X-100 in PBS and stained using alexa fluor 488 phalloidin (1:40) (Invitrogen) for 90 minutes followed by 4’,6-diamidino-2-phenylindole (DAPI) (1:500) (Sigma-Aldrich) for 10 minutes. The gels were washed with 1xPBS before 300 μm z-stacks were acquired on a Nikon-C2 laser scanning confocal microscope with a 20x air objective. The z-stacks were imported into IMARIS to generate surface renders, and the occupied volume.

Another set of gels were quantified using the PrestoBlue assay (Thermo Fisher Scientific) per manufacturer’s instructions to determine relative proliferation at day 2. The media in each well was replaced with a solution of 15 μL of the PrestoBlue reagent mixed with 135 μL of media. After 1 hours, 100 μL from each well were transferred into a 96-well plate and fluorescence was read using a BioTek plate reader at an excitation wavelength of 560 nm and an emission wavelength of 590 nm.

### Synthesis of PEG-3Tet-Alexa555

Polyethylene glycol-tri-tetrazine-Alexa Fluor 555 (PEG-3Tet-Alexa555) was synthesized through two base-catalyzed thiol-Michael additions in parallel by dissolving 1 mg of tetra-PEG-SH (MW: 20,000 Da) (NOF America, White Plains, NY) and 0.1 mg of methyltetrazine-PEG4-maleimide (MW: 514.53 Da) (Kerafast, Boston, MA) (Maleimide/SH ratio of 0.75) and 0.13 mg Alexa Fluor 555 C2 maleimide (MW: 1250 Da) (Thermo Fisher Scientific) (Maleimide/SH ratio of 0.25) in 0.5 mL CDCl3, then adding 0.1 µL of triethylamine (TEA) (3.5 molar equivalent). The mixture was stirred at room temperature for 4 hrs. The product was precipitated in 50 mL cold diethyl ether.

### Analysis of PEG-3Tet-Alexa555 Diffusion into HMP

MAP scaffolds containing PEG-3Tet-Alexa555 were assembled via inverse electron demand Diels-Alder tetrazine-norbornene click reaction, in which excess norbornene groups on the HMP were linked by a tetra-PEG-Tet solution containing 0.01 mM PEG-3Tet-Alexa555 to form Tet-MAP scaffolds with a labeled annealing location within each bead. (**Figure S5**) The final tetra-PEG-Tet annealing concentrations: 0.10 mM, 0.22 mM, 0.44 mM, and 0.89 mM for tetrazine/HA-Norb (Tet/HA) ratio of 2.5, 5, 10, and 20. The volume of tetra-PEG-Tet and PEG-3Tet-Alexa555 added to the HMP was consistent across the four Tet/HA conditions, 5/6 of the final Tet-MAP scaffold volume was HMP and 1/6 of the volume was 97 (v/v)% concentrated tetra-PEG-Tet (0.64 mM, 1.33 mM, 2.71 mM, and 5.46 mM, respectively) and 3 (v/v)% PEG-3Tet-Alexa555 (2.26 mM). Tet-MAP scaffolds with labeled annealing location were formed by combining and mixing HMP with concentrated tetra-PEG-Tet and PEG-3Tet-Alexa555 (PEG-Tet555) at the desired Tet/HA ratio and incubated for 1 hour at 37°C. All Tet-MAP scaffolds were formed using the 60 - 100 µm bead size filter range.

The distance that PEG-Tet555 diffused into a µbead was calculate in ImageJ after the center of a µbead from each Tet/HA condition was imaged using Nikon-C2 laser scanning confocal microscope with a 10x air objective. In ImageJ the gray values were measured across the diameter of each µbead analyzed. Using the center region, the average gray value of the background signal was determined. The diffusion distance into each µbead was the average distance that the average gray value of the background signal first intersected the gray value data from each edge. This was quantified for three HMP from each Tet/HA annealing ratio.

### Live/Dead

HDFs were encapsulated as described above, exchanging media every 2-3 days. Post-encapsulation a set of gels were used to quantify cell viability on day 1 and day 8. First, the live/dead kit (Invitrogen) solution was made per manufacture specification, 0.5 µL of calcein AM, 2 µL of ethidium homodimer-1, and 997.5 µL of media were combined. Then, the media in each well was replaced with 150 µL of the live/dead kit solution. After 20 minutes each well was replaced with media and a 300 µm z-stack was acquired using a Nikon-C2 laser scanning confocal microscope with a 10x air objective. These images were then analyzed using the 3D object counter toolkit in ImageJ to obtain the total number of live and dead cells for the 2.5 and 20 Tet/HA conditions. This data was then plotted using the fraction of total in Prism (GraphPad) generating the 95% confidence interval. (**Figure S6**)

---

**Figure S 1.**
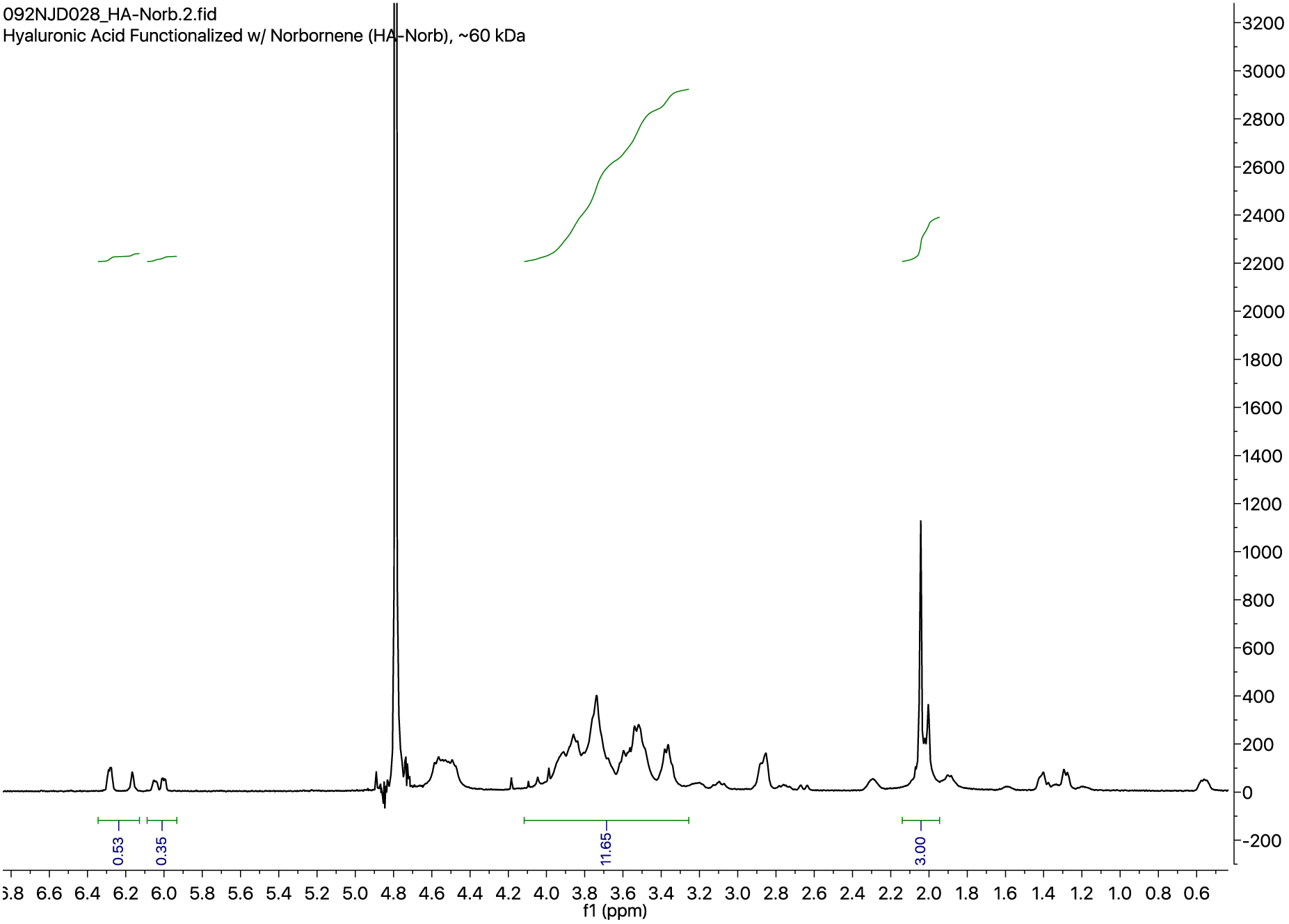
^1^H NMR of HA-Norb

**Figure S 2.**
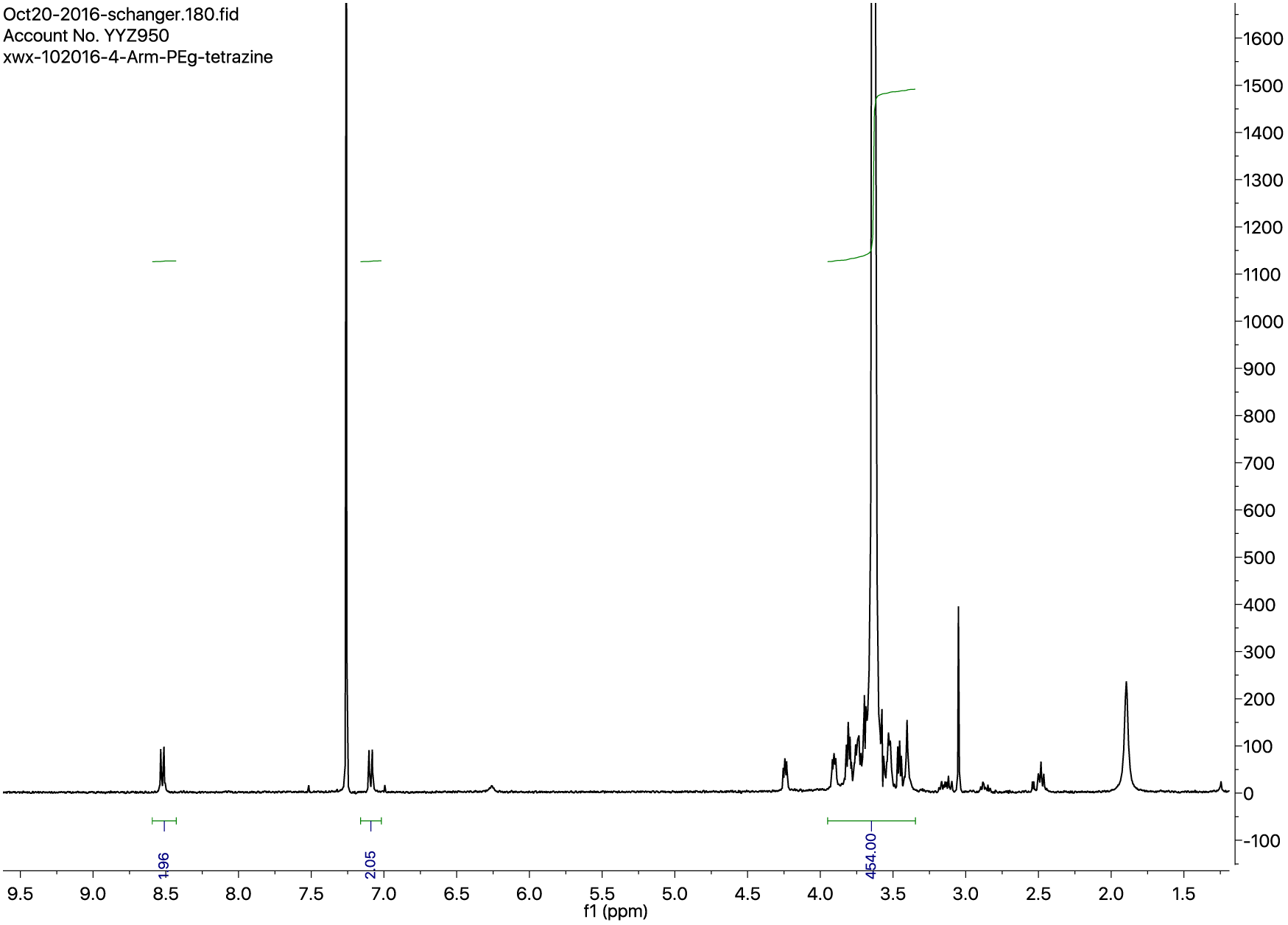
NMR of Tetra-PEG-Tet.

**Figure S 3.**
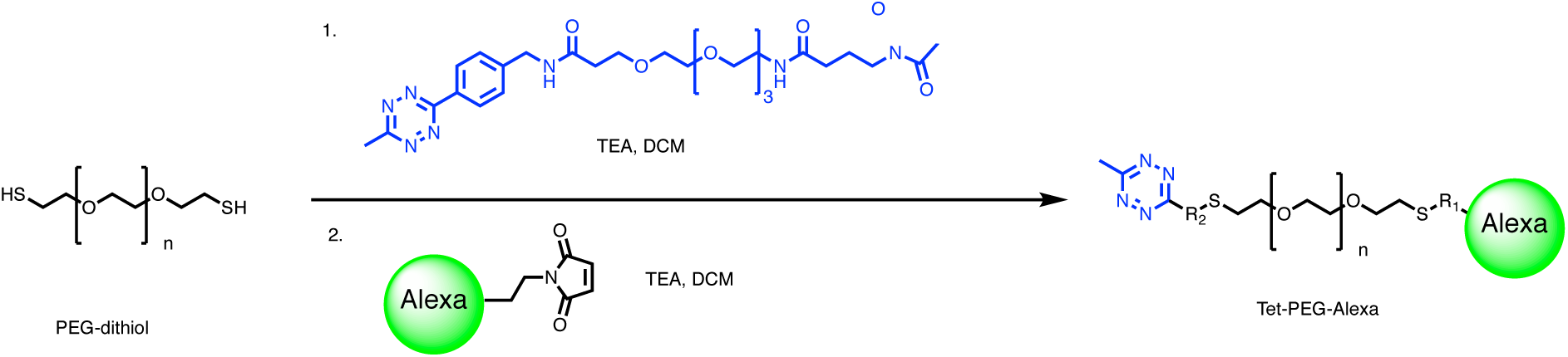
Synthesis scheme of Tet-PEG-Alexa

**Figure S 4.**
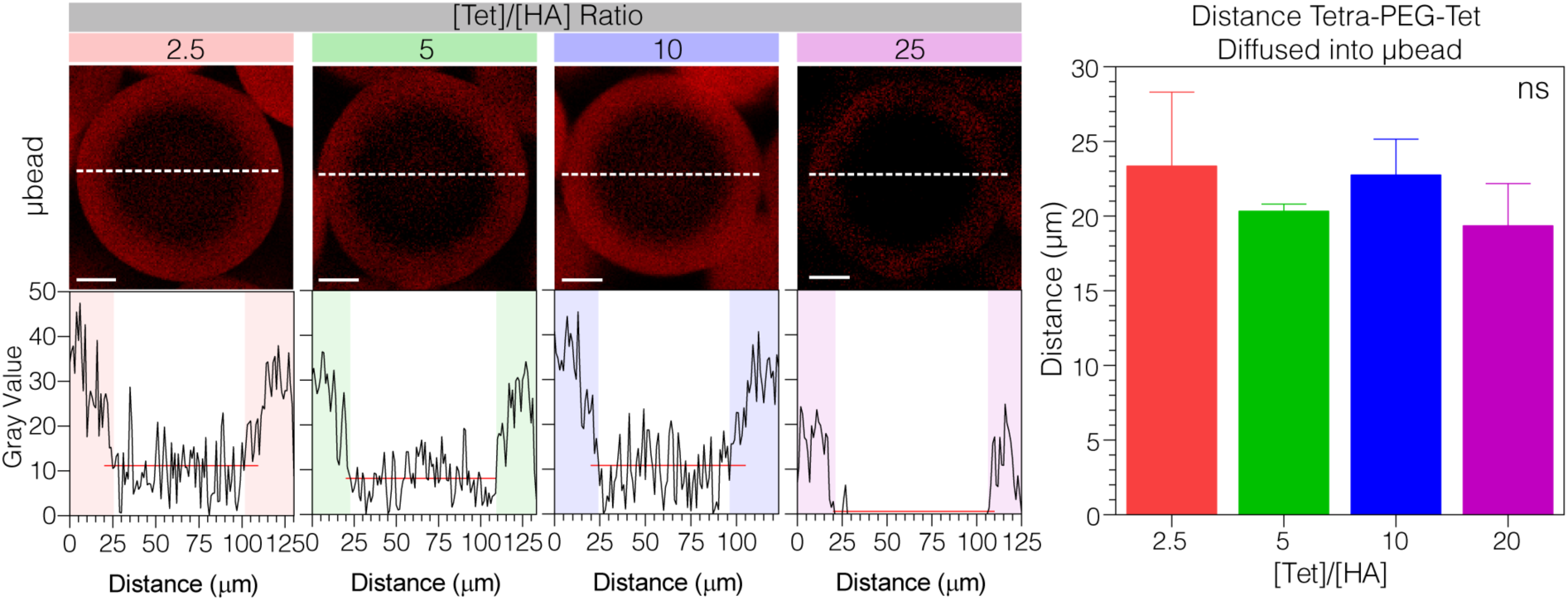
Diffusion analysis of Alexafluor-555-modified Tetra-PEG-Tet into individual HMP during annealing. (A) Representative images of HMP with each Tet/HA ratio are shown. White dashed lines represent the intensity plot path. The gray value for each bead is plotted along the path; the red line depicts the average background signal and scale bars = 10 µm. (B) The distance tetra-PEG-Tet diffused into individual HMP is plotted for each Tet/HA condition. Tukey’s multiple comparison test (P < 0.05). (n = 3 unless specified otherwise

